# Transcriptome dynamics in developing leaves from C_3_ and C_4_ *Flaveria* species reveal determinants of Kranz anatomy

**DOI:** 10.1101/473181

**Authors:** Kumari Billakurthi, Thomas J. Wrobel, Andrea Bräutigam, Andreas P.M. Weber, Peter Westhoff, Udo Gowik

## Abstract

C_4_ species have evolved more than 60 times independently from C_3_ ancestors. This multiple and parallel evolution of the complex C_4_ trait indicates common underlying evolutionary mechanisms that might be identified by comparative analysis of closely related C_3_ and C_4_ species. Efficient C_4_ function depends on a distinctive leaf anatomy that is characterized by enlarged, chloroplast rich bundle sheath cells and a narrow vein spacing. To elucidate molecular mechanisms generating this so called Kranz anatomy, we analyzed a developmental series of leaves from the C_4_ plant *Flaveria bidentis* and the closely related C_3_ species *Flaveria robusta* using leaf clearing and whole transcriptome sequencing. Applying non-negative matrix factorization on the data identified four different zones with distinct transcriptome patterns in growing leaves of both species. Comparing these transcriptome patterns revealed an important role of auxin metabolism and especially auxin homeostasis for establishing the high vein density typical for C_4_ leaves.

## Introduction

C_4_ plants evolved more than 60 times independently from their C_3_ progenitors (Sage et al., 2011; Sage, 2016). C_4_ photosynthesis increases the CO_2_ concentrations in the vicinity of Ribulose-1,5-bisphosphate carboxylase/oxygenase (Rubisco) up to 15-fold. This strongly suppresses the oxygenation reaction of Rubisco and accordingly photorespiration (Sage et al., 2012). Under current ambient CO_2_ concentrations (405 ppm) at 25°C, photorespiration is estimated to decrease the yield of soybean or wheat in the US by 36% and 20%, respectively (Walker et al., 2016). Environmental constrains such as high temperatures and drought further increases Rubisco oxygenase activity (Laing et al., 1974; Jordan and Ogren, 1984; Brooks and Farquhar, 1985; Parry et al., 2007).

In most C_4_ plants CO_2_ fixation is compartmentalized between two cell types, the bundle sheath (BS) and the mesophyll (M) cells. In the mesophyll phosphoenolpyruvate (PEP) is carboxylated by phosphoenolpyruvate carboxylase (PEPC), resulting in the 4-carbon compound oxaloacetate (OAA). OAA is converted to malate and/or aspartate, which is then transferred to the bundle sheath. Here the 4-carbon compounds are decarboxylated and the released CO_2_ is assimilated in the Calvin-Benson-Bassham cycle (CBB). The resulting pyruvate is transferred back to the mesophyll where the primary CO_2_ acceptor PEP is regenerated by pyruvate orthophosphate dikinase (PPDK) (Hatch, 1987).

C_4_ photosynthesis requires a particular leaf anatomy. As BS and M cells operate as a photosynthetic unit, direct contact of both cell types is necessary to ensure efficient photosynthesis. The BS is composed of the cells directly adjacent to the vasculature. Therefore, leaves of most C_4_ species exhibit high vein densities with a characteristic pattern in which two veins, each surrounded by BS cells, are separated by only two layers of M cells in a vein-bundle sheath-mesophyll-mesophyll-bundle sheath-vein layout. Bundle sheath cells of C_4_ plants often appear larger in cross-section compared to C_3_ species and contain more chloroplasts. This characteristic tissue pattern is also called Kranz anatomy (Haberlandt, 1904).

Current concepts hold that changes in leaf anatomy, leading to higher vein density and enlarged, chloroplast rich bundle sheath cells, are among the earliest steps of C_4_ evolution (Christin et al., 2013; Lundgren et al., 2014; Christin and Osborne, 2014; Bräutigam and Gowik, 2016; Sage et al., 2012). Several recent studies aimed at determining the factors governing the different developmental programs in C_3_ and C_4_ leaf development using comparative transcriptomics. They either analyzed the ontogeny of whole leaves (Wang et al., 2013a; Külahoglu et al., 2014) or sections of leaves that covered different stages of development (Aubry et al., 2014; Kümpers et al., 2017). Combined with in-depth analyses of several candidate genes, these approaches provided some insights into the changes of leaf development during C_4_ evolution as reviewed recently in (Sedelnikova et al., 2018) and (Kumar and Kellogg, 2018).

It became obvious that the leaf bundle sheath is equivalent to the root endodermis and the starch sheath of hypocotyls (Wysocka-Diller et al., 2000; Lim et al., 2005; Slewinski, 2013). The mechanisms controlling the development and differentiation of root endodermis, namely the short root/scarecrow regulatory system, are at least partially conserved in bundle sheath development and differentiation (Fouracre et al., 2014). It could be shown that *SCARECROW* and *SHORT-ROOT* mutants of maize as well as *Arabidopsis thaliana* exhibit distorted bundle sheath development (Slewinski et al., 2012, 2014; Cui et al., 2014). The overexpression of the maize *SCARECROW* gene (*ZmSCR1*) in Kitaake rice, on the other hand, did not lead to remarkable changes in leaf anatomy (Wang et al., 2017a).

Several transcription factors related to chloroplast development were identified, *e.g*., the GOLDEN2-LIKE (GLK) proteins (Hall et al., 1998; Fitter et al., 2002; Waters et al., 2009; Wang et al., 2013b) or B-class GATA transcription factors (Chiang et al., 2012; Hudson et al., 2013), mainly by their mutant phenotypes although they also showed up with several transcriptome centric approaches. Constitutive expression of one of the maize *GOLDEN2-LIKE* genes in rice induced chloroplast and mitochondrial proliferation in bundle sheath cells. The organelles increased in size and photosynthetic enzymes were induced mimicking the situation in C_4_ leaves (Wang et al., 2017b). So far, these *GLK* genes are the only known genes that can shift one of the signature anatomical traits towards the C_4_ state when solely overexpressed in a C_3_ leaf.

In leaves polar auxin transport and mesophyll differentiation are central to the development of a functional vascular network and control its structure and density. According to the widely accepted auxin canalization model (Sachs, 1969; Mitchison, 1981; Rolland-Lagan and Prusinkiewicz, 2005), vein differentiation is initiated by elevated auxin concentration due to polar auxin transport along strands of undifferentiated ground cells and terminate when photosynthetically active cells differentiate (Sud and Dengler, 2000; Scarpella et al., 2004; McKown and Dengler, 2010; Andriankaja et al., 2012). Differentiation of photosynthetic machinery is also associated with the exit from the phase of proliferation during leaf development (Adriankaja et al. 2012). Hence, venation density appears to be a function of auxin-related processes and the time spent in a differentiable state. In other words auxin x time = vein density. Külahoglu et al., (2014) observed that the differentiation of mesophyll cells is delayed in the leaves of the C_4_ species *Gynandropsis gynandra* compared to that of the closely related C_3_ species *Tarenaya hassleriana*. This prompted the hypothesis that more veins were initiated in the C_4_ leaf by prolonging their differentiation-competent state.

In the present study we performed an integrative analysis of the anatomy and the transcriptomes of series of developing leaves from the closely related C_4_ and C_3_ species *Flaveria bidentis* and *Flaveria robusta*. McKown and Dengler (2009) analyzed leaf development in these two species in great detail and showed that increased vein density in the C_4_ species is caused by formation of an additional minor vein order. In contrast to Kümpers et al., (2017) who followed leaf development in C_3_ and C_4_ *Flaveria* species by dissecting leaves of one stage into six pieces, we used a series of complete leaves of different ages. Nonnegative matrix factorization (NMF) of the transcript abundance data identified up to four highly distinct transcriptional patterns within the leaves of both species. Clearing of the leaves and determination of vein density mapped the different transcriptional programs to different developmental zones in the leaves.

## Results

### Transcriptomes of developmental leaf gradients from C_3_ and C_4_ *Flaveria* species

We performed RNA-seq on leaf developmental gradients from the C_4_ species *F. bidentis* and the C_3_ species *F. robusta*. The leaves analyzed cover the whole range of development from meristematic/primordial tissues (stage 0) to fully expanded leaves of about 10 cm length (stage 9) (Figure. 1A). RNA-seq analysis was performed on three independent biological replicates and generated on average 33 million reads per sample. Reads were mapped to a minimal set of coding sequences of the *Arabidopsis thaliana* transcriptome (http://www.arabidopsis.org/), as described by Bräutigam et al., (2011), with an average mapping efficiency of 53% (Supplemental Table 1). We were able to detect transcripts of 17,504 genes out of which 10,864 were expressed with at least ten RPKM (reads per kilobase of transcript, per million reads) in at least one sample and used for further analysis (Supplemental Table 2 and 3).

**Figure 1.**
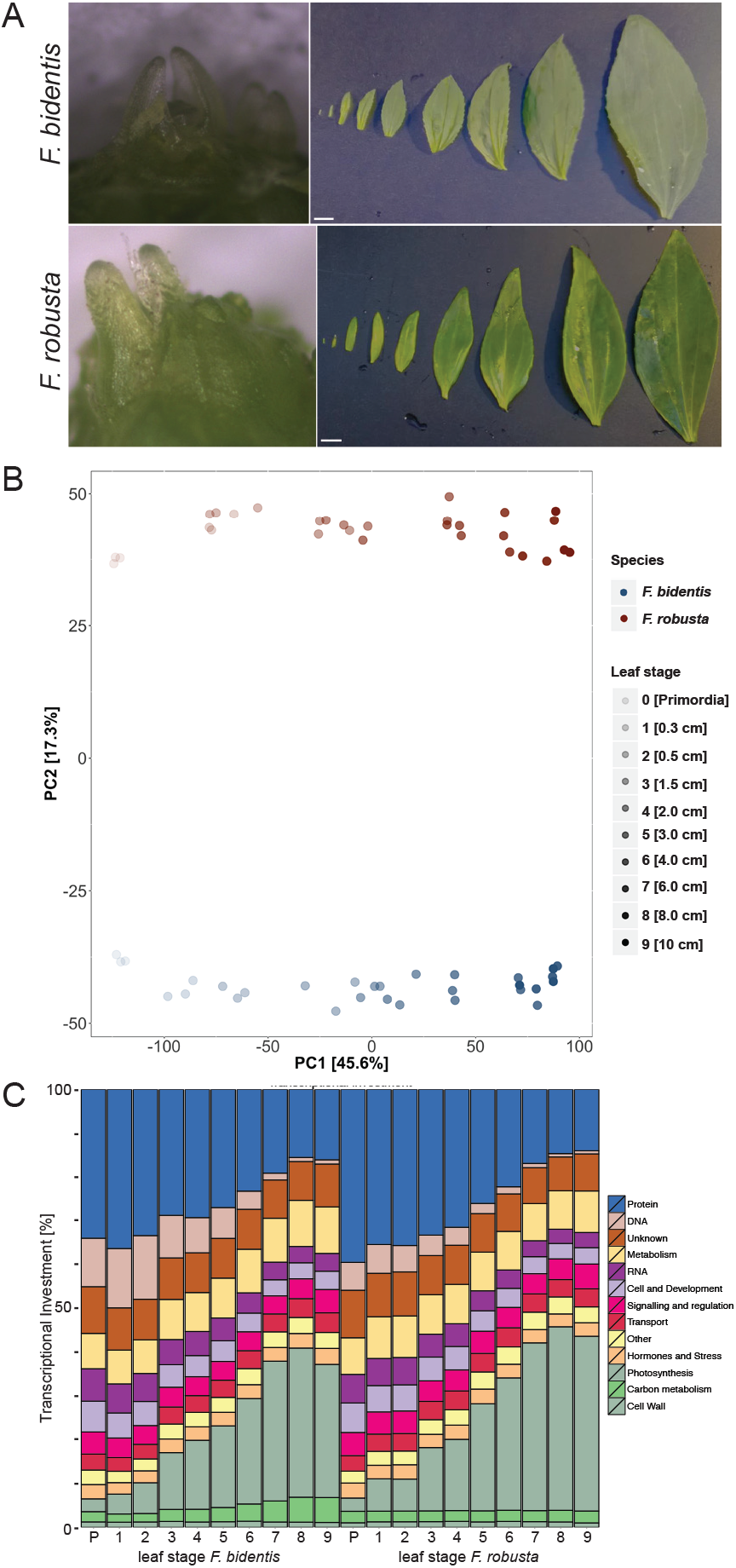
Leaf developmental transcriptomics of both C_3_ and C_4_ *Flaveria* species. A. Sampling of leafs from *F. bidentis* and *F. robusta* B. PCA of the developmental gradient for all leaf stages sequenced in this study. C. Transcriptional investment plot of the developmental gradient. The relative investment into different gene categories was calculated based on rpkm values.

Principal component analysis (PCA) of the dataset demonstrates high consistency between replicates (Figure 1B). The different leaf samples mainly vary by developmental stage of the leaves shown by a separation in principal component 1 explaining 45% of the total variation (Figure 1B). Species-specific variations including those related to the different types of photosynthesis are mainly represented by principal component 2 that explains 17% of the total variation (Figure. 1B).

Transcriptional investment, according to functional categories inferred from MapMan bins (Usadel et al., 2009) is largely in accordance with the PCA. Leaves from a comparable developmental stage show higher similarity between the different species than young and mature leaves of the same species (Figure 1C). Unlike the progression of expression in the C_3_ and C_4_ Cleomaceae species (Kühlaglu et al. 2014), the progression through leaf development exhibit near-identical timings for the C_3_ and the C_4_ the *Flaveria* species. While transcripts related to protein and DNA synthesis occupy a predominant role in young leaves, photosynthesis becomes dominant with progressing leaf age (Figure 1C). *Flaveria bidentis* exhibits a higher transcript investment into DNA synthesis in young tissues and into carbon metabolism in older leaves, while *F. robusta* has a slightly higher investment into photosynthesis.

### Changes of leaf anatomy and venation patterns in the two *Flaveria* species during development

To follow the phenotypic developmental progression of leaves from both species in our growth conditions, we performed cross sections within the upper quarter of the respective leaf for stages two to nine (Supplemental Figure 1). In both species anatomical changes during leaf differentiation progresses at a similar pace. While in stage two, leaves of both species mainly consist out of undifferentiated ground tissue, in stages three and four the first differentiating veins can be observed. A completely developed bundle sheath and palisade parenchyma is present at leaf stage five (3 cm, Supplemental Figure 1) in both species. Leaves of the C_4_ species *F. bidentis* are thinner, with fewer cell layers in all stages and stop further vertical expansion earlier than their C_3_ counterpart, between stages six (4 cm) and seven (6 cm).

In order to gain a deeper understanding of the developing vein system, all leaf stages were cleared using TOMEI-I (Hasegawa et al., 2016) and photographed with four times magnification using a light microscope. The microscopic images were stitched together and present an overview on the process of vascularization (Figure 2A). We quantified the vascular density along the leaf lamina from the base to the tip in a stepwise manner for leaf stages one to nine and calculated the vein density as a function of the relative leaf area. We could observe three clearly distinct areas that were not necessarily present in all different stages. Figure 2B shows the vascular densities of stage six leaves (4 cm in size) from both species that contain all different areas. The proximal leaf areas that are characterized mostly by meristematic cells exhibit a very low vein density in both species (Figure 2B) and represent the cell division zone of the leaves. This zone is followed by an area with a marked increase in vascular density indicating that the majority of veins differentiate in this area. The vein density in this area is higher in the C_4_ species (19.8 mm/cm^2^) compared to the C_3_ species (8.5 mm/cm^2^) (Figure 2B). Towards the tips of the older leaves from both species vein densities decreases to 11.2 mm/cm^2^ in *F. bidentis* and 5.7 mm/cm^2^ in *F. robusta* due to cell expansion (Figure 2B).

**Figure 2.**
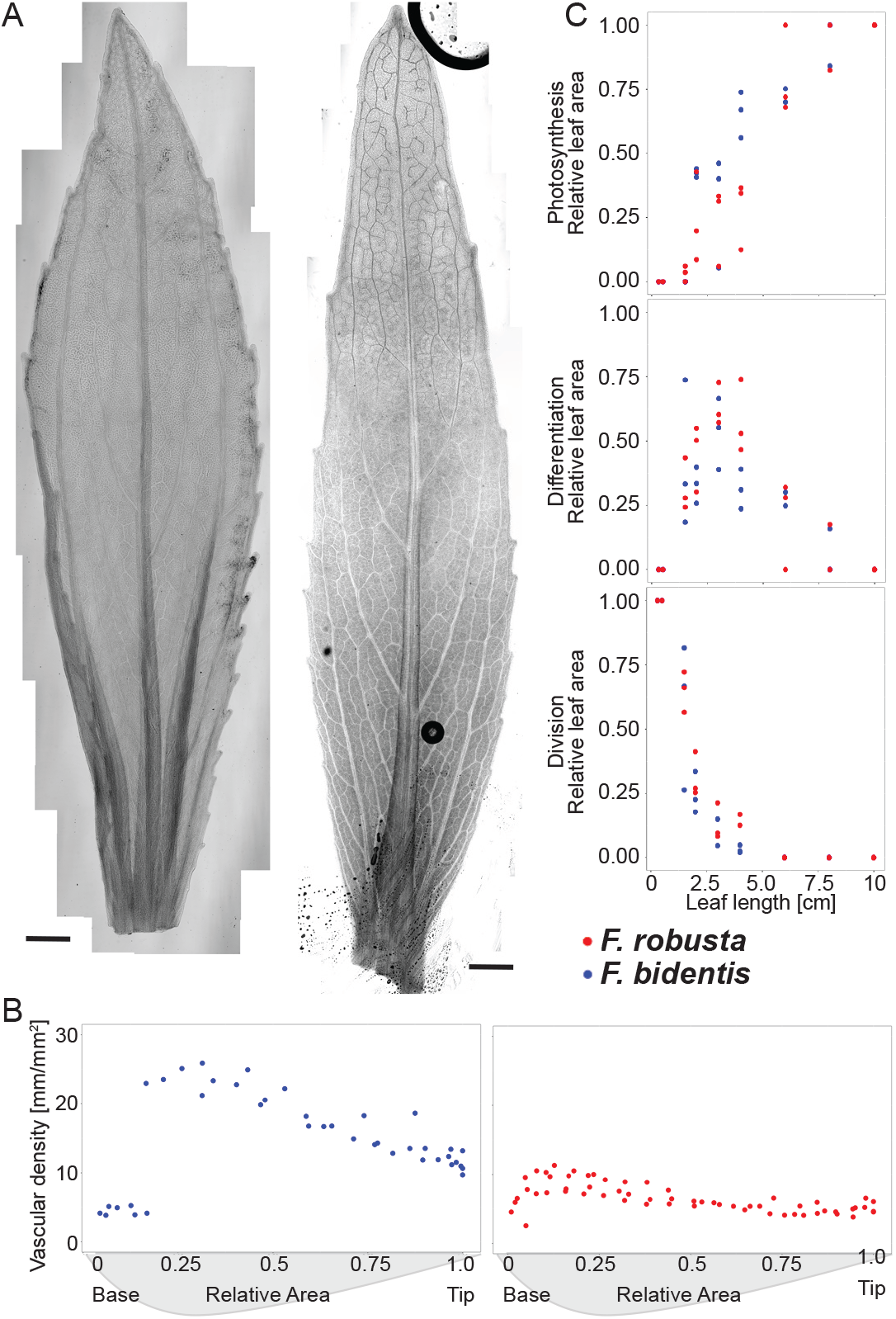
Anatomic changes during *Flaveria* leaf development. A. Stitched pictures of 2 cm leaf of *F. bidentis* (left) and *F. robusta* (right) that were used to determine the progression of vascular development. Scale bar = 1mm. B. Vascular density of 4 cm sized leaves presented as function of relative area. C. Relative proportions of Dividing (left), Differentiating (middle) and Photosynthetic (right) areas inferred from vascular density. *F. bidentis* in blue and *F. robusta* in red.

These characteristic vein densities were used to estimate the relative areas associated with cell division, cell/vein differentiation and mature, photosynthetic tissues. To this end differentiating tissues were considered to have vascular densities above 15 mm/cm^2^ in *F. bidentis* and 7 mm/cm^2^ in *F. robusta*. The area at the base of the leaves with lower vein density was categorized as consisting mainly out of dividing cells, whereas the distal area at the tips of the leaves were considered to consist out of mature and photosynthetic tissues. Based on this area estimates, cell division rapidly decreases in the first leaf stages reaching 0% of the leaf area in stage seven (6 cm). The majority of differentiation takes place between stage three (1.5 cm) and seven (6 cm), peaking at stage five (3 cm) in both species (Figure 2C).

### Estimation of gene expression patterns in the different developmental areas

Leaves have been separated into different zones based on cell area and shape (Adrianjaka et al. 2011) or based on venation and cell shape (Figure 2C, Kümpers et al. Kühlahoglu et al.). We sought to ab initio identify leaf zones based on deconvolution of transcript abundance patterns, independent of phenotypic observation. To this end, we consider the leaf as a composite of tissues with different developmental states, likely represented by the areas with different vein patterning. The proportions of the areas change depending on the developmental state of the leaf. When we assume that the expression of individual genes stays constant in the respective area of all leaf stages, the expression (e) of each gene (G) within a leaf sample (S) can be written as the sum of products of its area specific expression (e(A,G)) multiplied by the relative abundance of the area in a sample (A(S,A)) (Form.1A). In a sample, the relative abundances of the areas are identical for all genes, while the individual expression of all genes are area specific. This allows the formalization as matrix multiplication (Form.1B). As we consider relative tissue abundances, their sum has to add up to one. Using these constrains we applied Nonnegative matrix factorization (NMF) (Lee and Seung, 1999; Brunet et al., 2004) to deconvolute the gene expression data from all developmental stages (one to nine) from both species for all genes. We tested four algorithms and three different seeding methods for two to ten areas. To estimate the quality of the different approaches and the number of developmental areas we used the residual sum of squares (rss) and the average regression coefficient (R^2^) between the approximated and real values, performing NMF fifty times (Supplemental Figure 2).

The Lee and Seung, Brunet and KL algorithms yielded comparable results regarding the rss. NsNMF results varied depending on the enforced sparsity, with highly sparse solutions (theta 0.9) having high rss values (Supplemental Figure 2A). The seeding method on the other hand had no major impact on the resulting factorization (Supplemental Figure 2B). To further investigate the quality of the factorizations we compared their R^2^ values depending on the factorization rank (Supplemental Figure 2D). Consistent with the rss the average R^2^ of the nsNMF factorizations is highly dependent on sparsity, with high sparsities creating low R^2^ values. The highest R^2^ values were reached by the KL and Brunet algorithms, while the Lee and Seung method yielded slightly worse factorizations (Supplemental Figure 2D). The KL and Brunet algorithms have high identical R^2^ values at the proposed optimal factorization rank of five of 0.86 in *F. bidentis* and 0.79 in *F. robusta*. The Lee and Seung algorithm on the other hand yields R^2^ values of 0.64 in *F. robusta* and 0.79 in *F. bidentis* at the optimal factorization ranks of three in *F. robusta* and four in *F. bidentis*. Taken together the KL algorithm was comparable with Brunet and outperformed the others. KL was used with random seeding to perform the factorization for each species separately 500 times.

For both species, KL determined five areas as the ideal factorization when primordial tissues were included. In both species the primordial samples occupy a special position within the dataset (Supplemental Figure 3). Their presence creates an area that contributes to the majority of transcription in the primordial sample while this transcription pattern does not reoccur within the remaining leaf stages. k-means clustering hyphenated with GO enrichment analysis shows the genes peaking in this area to be enriched in hormone response, histone modification, floral organ development, meristem development and meiotic cell cycle (Supplemental Table 4, Supplemental Figure 4). The Primordial samples represent a separate functional state that is only marginally if at all present in the remaining leaves of the developmental gradient. Therefore, we focused the analyses on leaf stages by omitting primordial samples for subsequent analyses and performed the final NMF with four areas.

We calculated the quality of the fit provided from the NMF, area abundances and the area specific expressions for each gene via the regression coefficient (R^2^). Overall an average R^2^ of 0.81 was achieved for the *F. bidentis* data and an average R^2^ of 0.75 for the *F. robusta* data. This corresponds to 8690 of 10864 (80%) genes with a R^2^ > 0.7 in *F. bidentis* and 7558 of 10864 (70%) genes with a R2 > 0.7 in *F. robusta*. Modelled expression data together with the original read counts and correlation coefficients are provided in Supplemental Table 3.

While analysis of the anatomy points towards three different developmental areas (Figure 2C), prediction of different areas based on transcriptome data results in four different areas with quite different gene expression profiles (Figure 3A). Comparison of the predicted to the measured relative areas (Figure 3A and 2C, respectively) strongly suggests that the area with dividing cells is covered by the predicted area A1 according to high Pearson correlations of 0.97 for *F. bidentis* and 0.98 *for F. robusta*. Similar results were obtained for the differentiating area and the predicted area A2 with correlation coefficients of 0.92 and 0.95 for *F. bidentis* and *F. robusta*, respectively. The third measured area exhibits weaker correlations with either predicted area A3 or A4, exhibiting Pearson correlation coefficients of 0.61 and 0.76 for A3 and 0.84 and 0.75 for A4, in *F. bidentis* and *F. robusta* respectively. However, it highly correlates with the added value of the two predicted areas (0.98 *F. bidentis* and 0.95 *F. robusta*) suggesting that this anatomically apparently uniform area indeed consists of two different zones that can be distinguished by different gene expression profiles in both species.

**Figure 3.**
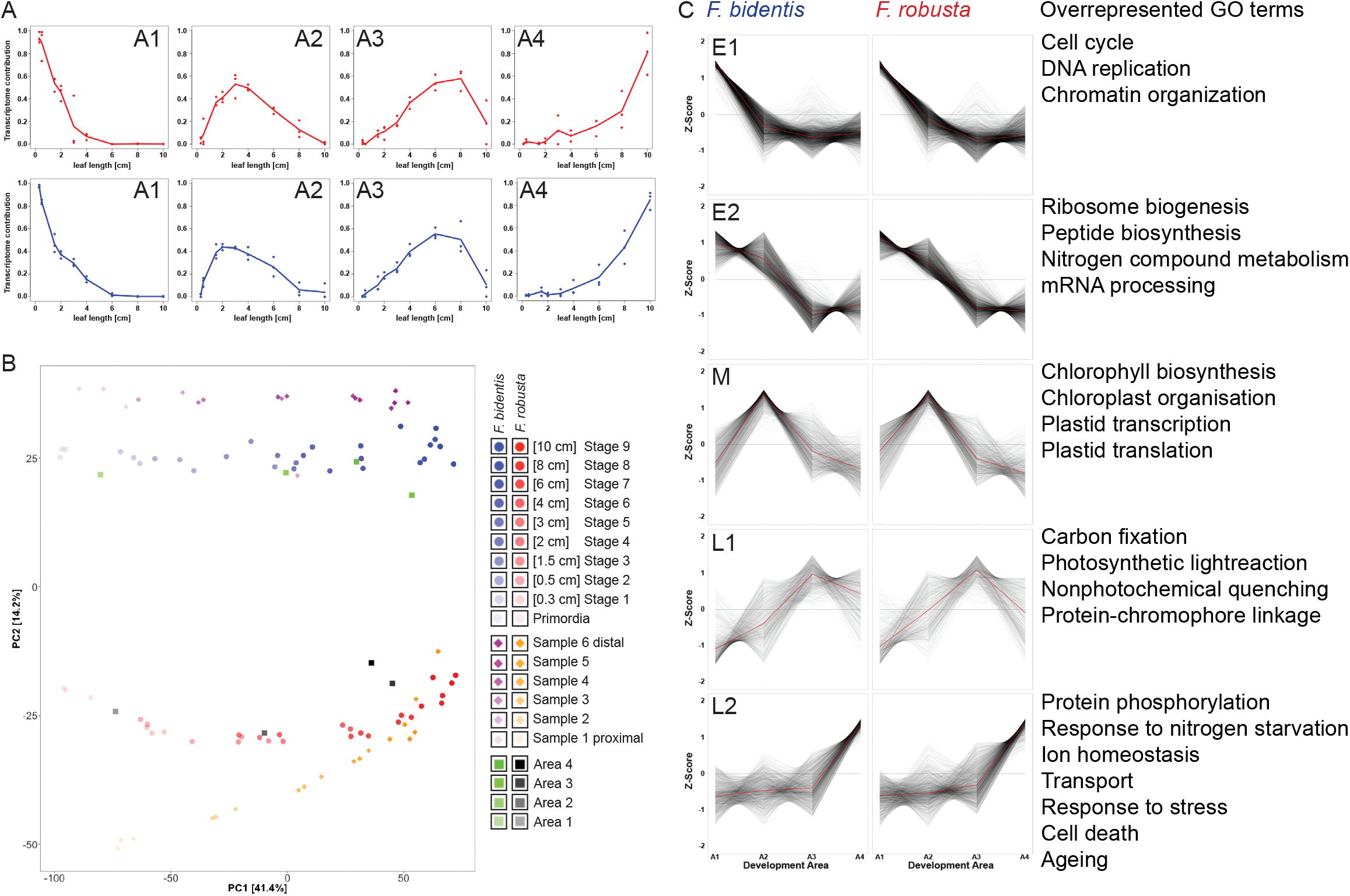
Unsupervised deconvolution of gene expression data. A. The relative contributions of each modelled area to the leaf transcriptomes in the developmental gradient. B. PCA of the developmental gradient for all leaf stages sequenced in this study (Düsseldorf) as well as the Flaveria slice transcriptomics from Kümpers *et al*. 2016. The modelled areas are depicted in green (*F. bidentis*) and black (*F. robusta*). C. k-means clustering based on the modelled gene coefficients. An excerpt of the overrepresented GO terms that are identical in both species is presented. A1-4 correspond to the Areas presented in A.

We compared the modelling results with sectional sequencing published by Kümpers et al., (2017), who sequenced the transcriptomes of 2 cm leaves from *F. bidentis* and *F. robusta* that were divided into six slices along the leaf from base to tip. The PCA of both datasets illustrates that they span a similar developmental space, with our modelled tissues representing central states within this progression for *F. robusta* (Figure 3B and Supplemental Figure 5) and slightly beyond for *F. bidentis*.

### Functional relevance of modelled leaf areas

To evaluate the biological significance of the predicted areas and modelled expression data, we applied k-means clustering to the deconvoluted dataset for both species separately. We obtained seven clusters for each species (Supplemental Figure 6) and GO enrichment analysis was performed using genes with a R^2^ score higher than 0.7 when comparing the modelled data to the real expression profiles in the analyzed species. Five clusters showed a remarkable expression pattern with an expression peak at the very early, early, middle, late and very late stage of the leaf development, respectively (Figure 3C, Supplemental Table 5). Accordingly, these clusters were termed E1, E2, M, L1 and L2. Area A1 is modelled to compose nearly 100% in young leaves and declines to 0% by leaf stage seven (6 cm) in both species (Figure 3A). Cluster E1 (Figure 3C) contains genes specifically up regulated within this first area (A1). A variety of GO categories are overrepresented in this cluster in both species. Most of them are characteristic for processes during early leaf development covering cell division, DNA replication, meristem initiation or the determination of bilateral symmetry (Figure 3C). For example, genes encoding CUP SHAPED COTYLEDON3 (CUC3), REVOLUTA (REV), YABBY3 (YAB3), components of the COP9 signalosome complex, histones, cyclins and cyclin dependent kinases (CDKs) are in cluster E1 in both species. Cluster E2 exhibits high expression in Areas A1 and A2 and is mainly characterized by ribosome biogenesis, peptide biosynthesis and protein biosynthesis.

Cluster M contains genes specific for area 2 (A2), contributing to 44% +/- 3% and 52.9% +/- 11% of the total transcriptome in *F. bidentis* and *F. robusta*, respectively. The area A2, peaks in leaf stage four (2cm) of *F. bidentis* and in leaf stage five (3 cm) of *F. robusta* (Figure 3A). Enriched GO terms in this cluster for both species are: Chloroplast organization, plastid transcription and translation as well as chlorophyll biosynthesis (Figure 3C).

None of the clusters was specific for area three (A3) although cluster L1 peaked in this area but also exhibit increased expression in area A4 in both species. GO enrichment in this cluster comprises photosynthetic light reactions and carbon fixation (Figure 3C). Cluster L2 peaks in area A4, which appears first at leaf stage five (3 cm) in both species and increases rapidly to contribute the majority of the leaf transcriptome in both species in the final stages (Figure 3A). GO terms enriched for this cluster are cell death, ageing, several transport processes and cellular response to starvation including the mobilization of nutrients (Figure 3C), which are all known to be related to the onset of senescence.

Based on these results we conclude that our deconvolution approach reflects leaf development as the modelled areas A1 to A4, according to their gene expression patterns, represent dividing tissues (A1), differentiating tissues with the onset of large-scale plastid development (A2), mature photosynthetic tissue (A3 and A4) and the onset of senescence (A4).

### Photosynthesis and photorespiration

Genes related to photosynthesis and photorespiration are mainly found in clusters L1 and L2 and are highly expressed in areas A3 and A4 (Figure 4 A; Supplemental Figure 7, 8 and 9). *F. bidentis* is known to be a NADP-ME C_4_ species and all landmark genes related to the C_4_ pathway in *Flaveria* (Gowik et al., 2011) like *CARBONIC ANHYDRASE* (*CA*), *PHOSPHOENOLPYRUVATE CARBOXYLASE* (*PEPC*), *NADP-MALATE DEHYDROGENASE* (*NADP-MDH*), *ASPARTATE AMINOTRANSFERASE* (*AspAT*), *NADP-MALIC ENZYME* (*NADP-ME*), *ALANINE AMINOTRANSFERASE* (*AlaAT*), *PYRUVATE, ORTHOPHOSPHATE DIKINASE* (*PPDK*), PPDK-regulatory protein (*PPDK-RP*) are highly expressed in areas A3 and A4 of leaves of the C_4_ species *F. bidentis* but not in leaves of the C_3_ species *F. robusta* (Supplemental Figure 7).

**Figure 4.**
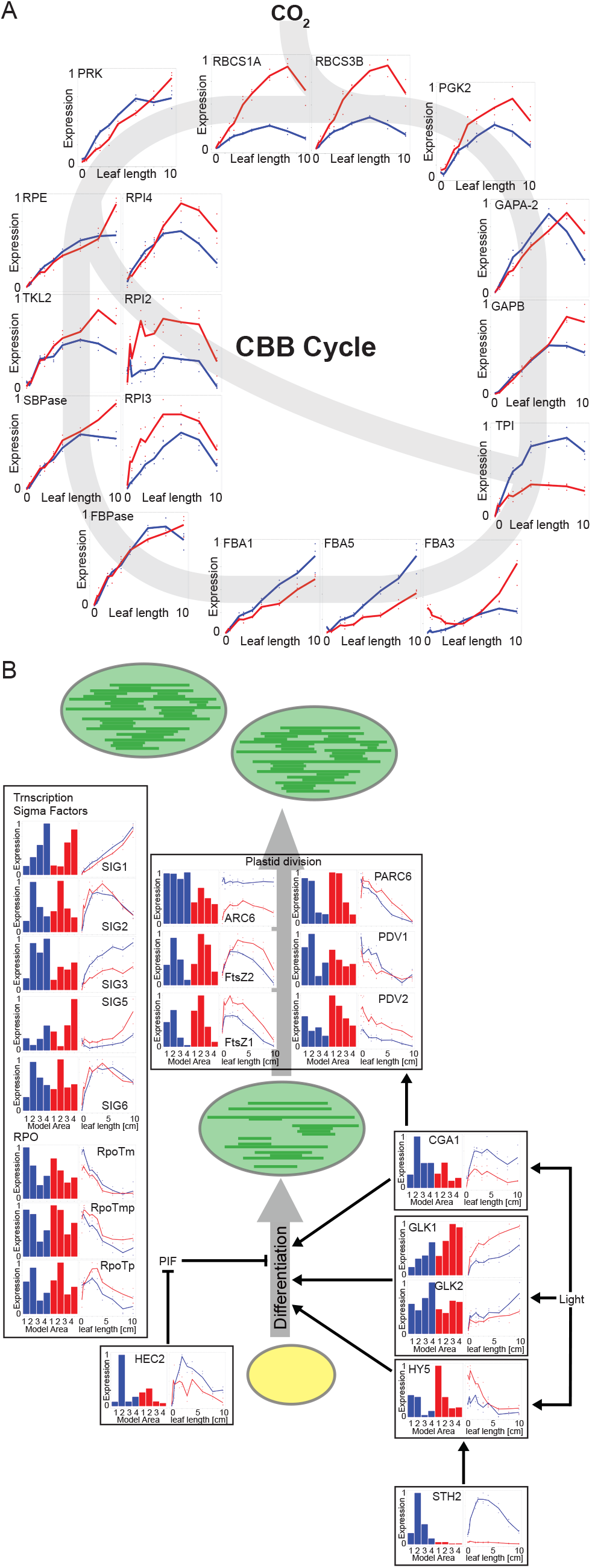
Expression patterns of the CCB and chloroplast assembly genes. A. Expression of Calvin Benson Bassham Cycle genes throughout the gradient, normalized to maximum expression per species. B. Expression of genes connected to Plastid differentiation and division. Modelled and original data is normalized to its maximum within each species.

Consistent with the results of Nakamura et al., (2013) transcripts related to both complexes involved in the cyclic electron transport, the NADH dehydrogenase-like (NDH) complex and PGR5/PGRL1 are more abundant in the C_4_ species while the expression of most transcripts peak in area A3 (Supplemental Figure 8). Genes encoding the proteins of the CBB cycle are most highly expressed in areas A3 and A4 of both species. Most of the CBB genes and especially the Rubisco small subunit genes are more highly expressed in the C_3_ species (Figure 4A). As to be expected, transcripts related to photorespiration are more abundant in the C_3_ species *F. robusta*. The expression of photorespiratory genes is highest in leaf areas A3 and A4 (Supplemental Figure 9). The expression of genes related to photosynthesis and photorespiration is in accordance with our modelling result, that areas A3 and A4 represent mature photosynthetic active tissues.

### Chloroplast development and division

Chloroplast differentiation is a key regulatory step in cell development to exit from proliferation stage (Andriankaja et al., 2012) and is associated with termination of vein formation (Scarpella et al. 2004). We have identified several transcription factors that positively regulate chloroplast development and/or biogenesis being upregulated during C_4_ leaf differentiation compared to C_3_ leaves (Figure 4B). In the light the ELONGATED HYPOCOTYL 5 (HY5) transcription factor stimulates photomorphogenesis (Waters and Langdale, 2009). It is known that B-class GATA transcription factors (Chiang et al., 2012; Hudson et al., 2013) and GOLDEN2-LIKE transcription factors (GLK1 and GLK2) (Hall et al., 1998; Fitter et al., 2002; Waters et al., 2009; Wang et al., 2013b) positively regulate the chloroplast development. The gene *CGA1* (ATGATA22; *CYTOKININ-RESPONSIVE GATA FACTOR* 1) is upregulated in the C_4_ species *F. bidentis*. In both species, *CGA1* attained maximum expression in leaf area A2 whereas transcript abundances are two-fold higher in area A2 of the C_4_ leaves (Figure 4B). CGA1 positively regulates chloroplast development and division (Chiang et al., 2012). Transcripts of *GLK-2* did not differ much between C_3_ and C_4_ *Flaveria* species while *GLK-1* levels are slightly up in *F. robusta* (Figure 4B). We have identified further transcription factors like STH2 or BBX21 (SALT TOLERANCE HOMOLOG 2 OR B-BOX CONTAINING PROTEIN 21) and HEC2 (HECTATE2) being upregulated in the C_4_ leaves. STH2/BBX21 and HEC2 indirectly promote photomorphogenesis by positively regulating the transcription of *HY5* and by inhibiting the expression of *PHYTOCHROME INTERACTING FACTOR1* (*PIF1*) respectively (Xu et al., 2016; Zhu et al., 2016) (Figure 4B). We hypothesize that up-regulation of these factors in the C_4_ species is related to chloroplast development in the bundle sheath, since overall chloroplast development seems to be quite similar in both species as indicated by genes related to plastid division and chloroplast gene expression (Figure 4B).

Plastid division mainly occurs in in area A2 in both species, as is reflected in the transcription patterns of *FtsZ1* and *FtsZ2* (Figure 4B). Chloroplast transcription peaks in a similar manner in area A2, with the highest predicted expression of the plastidic RNA polymerase (*RpoTp*) as well as *SIGMA* transcription factors, two (*SIG2*) and six (*SIG6*) (Figure 4B). The mitochondrial RNA polymerase gene (*RpoTm*) on the other hand shows the highest expression in area A1, while RopTmp believed to interact with both organelles (Kühn et al., 2007) exhibits similar levels in both areas (Figure 4B).

### Genes related to bundle sheath differentiation are upregulated in the C_4_ species

Bundle sheath tissue in leaves is equivalent to the root endodermis and the starch sheath of hypocotyls and the development of bundle sheath cells is at least partially conserved to root endodermis development and differentiation (Kyung Yoon et al., 2016; Wysocka-Diller et al., 2000; Lim et al., 2005; Slewinski, 2013; Cui et al., 2014). It is known that the interplay of the GRAS-type transcription factors SCARECROW (SCR) and SHORT-ROOT (SHR) is essential to specify the root endodermis identity as well as for the correct bundle sheath development in the C_3_ species *Arabidopsis* and the C_4_ species maize (Slewinski et al., 2012; Cui et al., 2014; Slewinski et al., 2014). In both *Flaveria* species, transcript abundances of genes encoding SCR and SHR attain a maximum in dividing tissue (area A1) and decline during further leaf development. Transcripts of both, *SCR* and *SHR* are upregulated in *F. bidentis* (C_4_) leaf areas A1 and A2 compared to the C_3_ species (Supplemental Figure 10). The activity of *SCR* and *SHR* is regulated by interactions with intermediate domain (IDD) transcription factors that also fulfil roles in several other developmental processes (Welch et al., 2007; Ogasawara et al., 2011; Coelho et al., 2018). Recently, IDDs were proposed to be also involved in bundle sheath development (Coelho et al., 2018). Transcripts of *IDD2, IDD5, IDD10* (*JKD*) and *IDD12* genes were specifically more abundant in differentiating tissue (area A2) of the C_4_ species, while *IDD15, IDD7, IDD16* and *MGP* abundance peaked in area A1 of both species. IDD15 and IDD16 were again higher in the C_4_ species, while MGP and IDD7 retained similar transcript levels (Supplemental Figure 10). Expression patterns of SCARECROW-LIKE gene family members (*SCL*) are more similar in both species. Only *SCL3* is upregulated in area 2 in C_4_ species *F. bidentis* (Supplemental Figure 10). *SCL3* promotes root endodermal cell elongation (Heo et al., 2011). Genes involved in bundle sheath development and differentiation can be expected to be upregulated in developing leaves of C_4_ species compared to C_3_ species since more bundle sheath cells have to develop due to the higher vein density in C_4_ leaves.

### The timing of mesophyll and bundle sheath differentiation is comparable in both species

Analysis of leaf development in C_3_ and C_4_ Cleomaceae species revealed that increased vein formation in the C_4_ species *Gynandropsis gynandra* is likely related to a delay in mesophyll cell differentiation during leaf development (Külahoglu et al., 2014). This does not seem to be the case during leaf development in C_3_ and C_4_ *Flaveria*. Leaf development proceeds quite similar in both species regarding to changes either in anatomy or in gene expression. Completely developed bundle sheath and palisade parenchyma appear in the same leaf stage in both species (stage 4; Supplemental Figure 1). Expression of photosynthetic genes and genes related to chloroplast development peak in the same leaf stages (Figure 3C and Figure 4). Overall transcriptional investment is quite similar in both species for all analyzed leaf stages (Figure 1A). When analyzing our modelled data, the predicted leaf areas are again very similar for both species (Figure 3A). Leaf area A1 is modelled to compose nearly 100% in young leaves and declines to 0% by leaf stage seven (6 cm) in both species. Area A2, peaks in leaf stage four (2cm) of *F. bidentis* and in leaf stage five (3cm) of *F. robusta* (Figure 3A) whereas area A3 peaks in leaf stages seven and eight of both species respectively. The leaf area A4 first appears in leaf stage five and increases in both species with leaf size (Figure 3A). These findings are comparable with the results of Kümpers et al., (2017) who also did not report any delays in leaf development or cell differentiation in C_4_ compared to C_3_ *Flaveria*. This suggest that in C_4_ *Flaveria* leaves, the auxin term of the equation auxin x time = vein density might be modified.

### Auxin metabolism is upregulated in dividing and differentiating tissues of the C_4_ species compared to the C_3_ species

In leaves polar auxin transport is central to the development of a functional vascular network and controls its structure and density. The local auxin maxima are created by auxin synthesis in dividing tissues and through transport facilitated by the concerted expression of different PIN proteins (Verna et al., 2015). Accordingly, genes related to the main Indole-3-acetic acid (IAA) synthesis pathway, the Indole-3-pyruvic acid (IPA) pathway (Mashiguchi et al., 2011), e.g. *TAR2* (*TAA RELATED PROTEIN 2*) and *YUCCA1* (*YUC1*) are mainly expressed in leaf area A1 and correspondingly leaf stages one and two of both species (Figure 5). YUC1, encoding the enzyme catalyzing the rate-limiting step of the IPA pathway (Mashiguchi et al., 2011) is higher expressed in the C_4_ species compared to the C_3_ species indicating that IAA synthesis might be enhanced. Additionally, AMIDASE 1 (AMI1) catalyzing the synthesis of IAA from Indole-3-acetamide is highly expressed in the differentiating area of both species and might contribute to auxin homeostasis.

**Figure 5.**
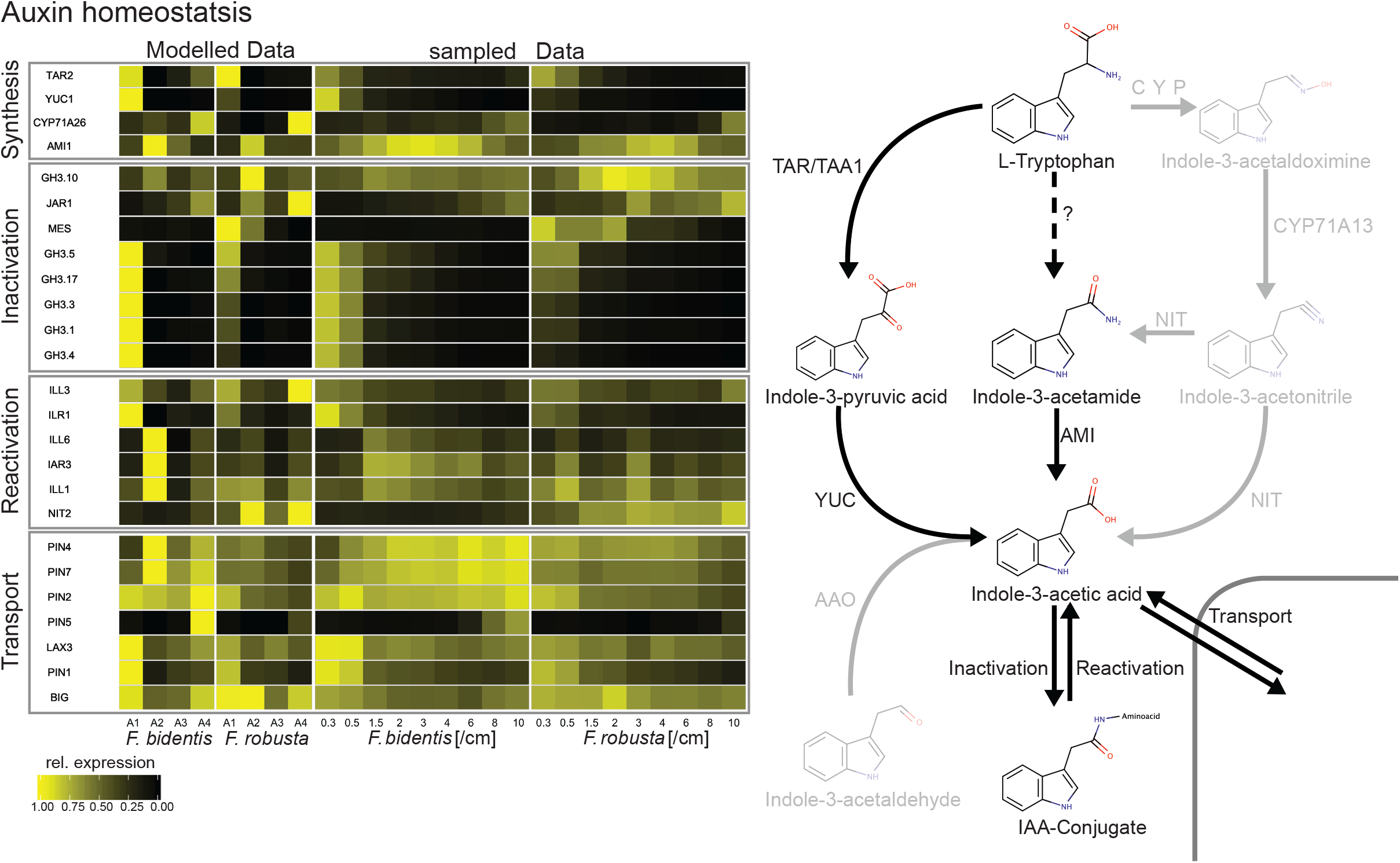
Heatmap of genes related to auxin synthesis and homeostasis. Gene expression is presented normalized separately for the modelled and the sampled data to the respective maximum in both species.

IAA can be inactivated by sequestration into amino acid conjugates (Nakazawa et al., 2001; Takase et al., 2004; Staswick et al., 2005). This reaction is catalyzed by Gretchen Hagen 3 (GH3) enzymes (Staswick et al., 2002, 2005). Genes encoding these enzymes are known to be induced by auxin (Hagen and Guilfoyle, 1985). Accordingly, we observe expression of *GH3* genes mainly in leaf area A1 in both species where auxin is synthesized. The majority of the *GH3* genes covered by our data set exhibit a marked increase in expression in the C_4_ species (Figure 5). IAA can be set free from amino acid conjugates and thus be reactivated by hydrolyzing enzymes (Bartel and Fink, 1995; Davies et al., 1999; Rampey et al., 2004). Genes encoding such enzymes like *ILL* (IAA-*AMINO ACID CONJUGATE HYDROLASE*) and *ILR* (*IAA-LEUCINE RESISTANT*) genes are more highly expressed in leaf area A1 and especially leaf area A2 of the C_4_ species compared to the C_3_ species (Figure 5).

Upregulation of auxin conjugating and reactivating genes in the division and differentiation areas of C_4_ leaves implies a higher capacity of auxin homeostasis in the C_4_ species *F. bidentis*. To test this hypothesis, we treated three-week-old C_3_ and C_4_ *Flaveria* plants for three weeks with the synthetic auxins NAA (1-Naphthaleneacetic acid) and 2,4-D (2,4-Dichlorophenoxyacetic acid) via leaf spraying. 2,4-D cannot be sequestered effectively into amino acid conjugates by GH3 proteins while NAA is a GH3 substrate (Staswick et al., 2005).

In both species 2,4-D application was lethal in concentration above 10 μM. At 10 μM it caused a severe phenotype with distorted leaf shape and significantly increased vascular density in both species (Figure 6A). With 1 μM 2,4-D leaf shape was still distorted in both species. On the tissue level the vascular structure of the C_4_ plant *F. bidentis* was highly perturbed with fused veins forming plate like structures at both concentrations and a significant increase in vein density at 10 μM 2,4-D (Figure 6B, C), while the C_3_ species showed an increase in vascular density at 1 μM 2,4-D (Figure 6C). NAA changed the overall leaf shape and significantly increased vascular density in *F. robusta* (C_3_) at its highest concentration of 790 μM but not in *F. bidentis* (C_4_) (Figure 6A, C). Similar effects, specifically an increased vein density, are reported when *A. thaliana* leaves accumulate auxin (Mattsson et al., 1999; Sieburth, 1999). These data support the hypothesis that auxin homeostasis in *F. bidentis* is altered compared to *F. robusta* as a result of increased sequestration of active auxin into inactive amino acid conjugates. Hence, the C_4_ species tolerates higher auxin levels than the C_3_ plant without clear alterations of leaf anatomy in case of the GH3 substrate NAA. 2,4-D generates effects at lower concentrations compared to NAA since it cannot be sequestered effectively into amino acid conjugates by GH3 proteins while NAA is a GH3 substrate (Staswick et al., 2005).

**Figure 6.**
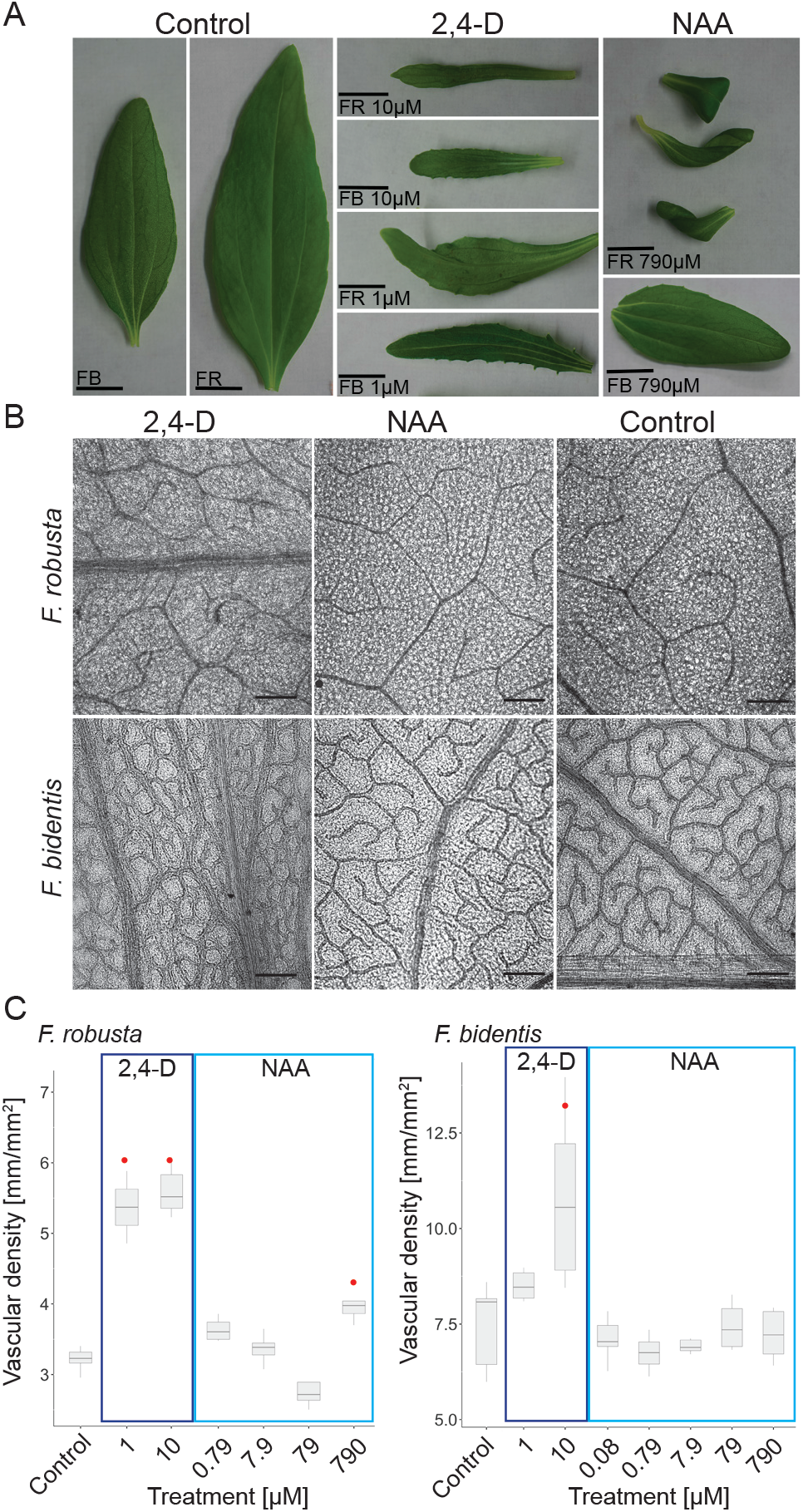
Anatomical changes of *F. bidentis* and *F. robusta* leaves sprayed with synthetic auxins. A. Leaf shapes of plants sprayed with 2,4-D, NAA or the 0.1% DMSO control (Scale bar – 1 cm). B. Vascular structure of plants sprayed with 10 μM 2,4-D, 700 μM NAA or the 0.1% DMSO control (Scale bar – 250 μM). C. Vascular density of plants sprayed with varying concentrations of 2,4-D, NAA and the 0.1% DMSO control.

Genes related to auxin transport are also differentially expressed in both species. While PIN1 expression is highest in dividing tissues (area A1) and decreases rapidly during differentiation (area A2) in both species its expression is higher in the *F. bidentis* (C_4_). PIN7 and PIN4 expression peak during differentiation (area A2) and PIN2 and PIN5 expression peak in area A4 of the C_4_ leaves while these genes exhibit much lower transcript levels in the C_3_ leaves (Figure 5).

### Genes related to auxin signaling and auxin induced genes are differentially expressed in C_3_ and C_4_ leaves

Genes encoding the auxin receptors TIR1 (TRANSPORT INHIBITOR RESPONSE 1), AFB1 and AFB3 (AUXIN SIGNALING F-BOX 1 and 3) are specifically upregulated in leaf area A2 of the C_4_ species *F. bidentis* (Supplemental Figure 11) whereas their expression is more uniform but lower in areas A1 and A2 of the *F. robusta* (C_3_) leaf (Supplemental Figure 11). Expression of other, more downstream, components of auxin signalling e.g. *IAA* and *ARF* genes appear to be more similar in both species (Supplemental Figure 11). Although we found that expression patterns of *IAA5* and *IAA6* as well as of *ARF3, ARF19* and *ARF16* show peaks in leaf area A2 in the C_4_ species that are missing in the C_3_ species (Supplemental Figure 11).

In accordance with the expression patterns of genes related to auxin synthesis, homeostasis and transport we observed an area specific induction of genes known to be regulated by auxin, like the SAUR gene family (*SMALL AUXIN REGULATED RNAs*) (Supplemental Figure 12). The majority of these genes were most highly expressed in leaf area A1 of both species. Although we identified several genes like *SAUR10, 12, 16, 53* or the auxin-induced dormancy related gene AT1G54070 that were exclusively or most highly expressed in leaf area A2 of the C_4_ species *F. bidentis* (Supplemental Figure 12). This finding further supports the hypothesis that increased expression of genes related to auxin synthesis and homeostasis in leaf areas A1 and A2 of *F. bidentis* translate to higher auxin levels especially in leaf area A2 of the C_4_ species compared to the C_3_ species.

## Discussion

C_4_ plants evolved more than 60 times independently from C_3_ ancestors. Since C_4_ photosynthesis demands Kranz anatomy for its function, leaf anatomy was subject to change during most of these independent evolutionary events. Kranz anatomy is characterized by high vein density and enlarged bundle sheath cells with elevated numbers of organelles, especially chloroplasts. To unravel the molecular mechanisms underpinning altered leaf anatomy in C_4_, we comparatively analyzed anatomy and transcriptomes during leaf ontogeny in the closely related C_3_ and C_4_ species *F. robusta* and *F. bidentis*. Analyses of vein densities in a developmental series of leaves from both species point towards the existence of three clearly distinguishable zones in leaves of both species. A cell division zone, a cell differentiation zone and a zone with mature, photosynthetic cells. We used nonnegative matrix factorization to deconvolute transcriptome data from whole leaves of sequential stages covering the growth process in both species. This revealed the existence of four highly distinct transcriptional patterns within the leaves of both species.

### NMF as a tool for the deconstruction of developmental processes

Nonnegative matrix factorization was introduced as a tool to recognize patterns for the decomposition of images into meaningful features (Lee and Seung, 1999), spectral data analysis or denoising of audio signals. In biology, it has been applied to a variety of datasets ranging from EEG based imaging (Lu and Yin, 2015; Delis et al., 2016) and muscle electrographs to transcriptomic datasets from microarrays or RNA-Seq experiments (Kong et al., 2011; Zhang et al., 2016; Shao and Höfer, 2017; Duren et al., 2018; Ebied et al., 2018). Regarding transcriptome data, the initial idea was published by Brunet et al., (2004). They applied NMF on a gene by sample matrix to identify metagenes from microarray data of different tumor cells (Brunet et al., 2004). An alternative option is the use of NMF to factorize sample by gene matrices to create so called metasamples. In plants Wilson et al. (2012) used NMF to deconstruct the AtGenExpress datataset into the metagenes shaping the transcriptional landscape of *Arabidopsis thaliana*. Recently (Shao and Höfer, 2017), analyzed metasamples in single cell gene expression datasets and were able to demonstrate that these can be used to correctly classify cell types and detect their signature genes.

In our work, we restricted the metagenes to reflect the relative composition of different stages of leaf development. To this end metagenes were normalized to a sum of 1 in every sample. The assumption, that the expression of individual genes stays constant in a respective developmental area throughout all the different leaves, is of course an oversimplification of the real situation and might be inaccurate for certain genes. Therefore, we critically assessed the results of the NMF. Using this method, we were able to obtain a realistic representation of the tissue composition of the different leaves ab initio from the transcriptome data in both species. The cell division zone as well as the cell differentiation zone, as defined by their anatomical features, could be clearly identified as indicated by the high correlations of modelled and measured area sizes (Figures 2 and 3). Additionally, the factorized data indicate that the area of mature, photosynthetic cells can be subdivided into two areas with clearly different transcript patterns. Clustering of gene expression data and analysis of enriched GO terms indicate that this is due to the onset of senescence in this area of older leaves. Similarly, maize leaves show senescence associated transcripts towards the tip of the leaf (Pick et al. 2011). The modelled expression data correctly capture the signature expression differences between both species as indicated by genes and metabolic pathways for which we have prior expectations, such as C_4_ photosynthesis, photorespiration, or the CBB cycle (Figure 4, Supplemental Figure 7, Supplemental Figure 8). Finally we compared the modelled expression data with the results of Kümpers et al., (2017) who analyzed base to tip maturation gradients of 2 cm long C_3_ and C_4_ *Flaveria* leaves. We found both datasets to be highly similar when compared via PCA (Figure 3, Supplemental Figure 5) except for the absence of area A4 in *F. bidentis* in the sectional data. Our results indicate that indeed 2 cm leaves are predicted to lack significant leaf area of the type A4 which is present only in larger, older leaves (Figure 3A).

### Leaf development proceeds very similar in both *Flaveria* species

Both, the analyses of the leaf anatomy as well as the analysis of the corresponding leaf transcriptomes indicate that leaf development in the C_4_ species *F. bidentis* is quite comparable to the leaf development in the C_3_ species *F. robusta*. Leaves of the same age have a comparable size and the distribution of anatomically distinguishable developmental zones are very similar in leaves of the same size from both species (Figure 2). As the main difference we identified the overall higher vein density in leaves of the C_4_ species.

Accordingly, the transcriptomes of leaves of the same stage are quite similar in both species. Transcriptional investment is very similar when leaves from the same age are compared and the PCA of all our RNA-Seq data indicates that the variation between the samples from different developmental leaf stages is up to three times higher than the species-specific variation (Figure 1B, 1C). Key developmental events like chloroplast division and differentiation occur simultaneously in both species. These findings are in line with earlier studies of leaf development in C_3_ and C_4_ *Flaveria* species (McKown and Dengler, 2009; Kümpers et al., 2017).

### Changes in auxin homeostasis could be related to higher vein density in the C_4_ *Flaveria*

The transcriptome analysis of developing leaves from a C_3_ and a C_4_ *Flaveria* species indicate that the young leaves of the C_4_ plants exhibit higher capacity for auxin synthesis and auxin homeostasis. Genes related to the main auxin synthesis pathway in dicot leaves (Mashiguchi et al., 2011) as well as genes related to the sequestration of auxin into amino acid conjugates, e.g. several *GH3* genes (Staswick et al., 2002, 2005), are more highly expressed in the cell division area of the C_4_ leaves than in the C_3_ leaves (Figure 5). On the other hand, we found genes related to re-activating the auxins due to hydrolase activity, e.g. *ILL, IAR* and *ILR* genes (Bartel and Fink, 1995; Davies et al., 1999; Rampey et al., 2004), up-regulated in the cell differentiation area of the C_4_ compared to the C_3_ leaves. This implies higher auxin availability in the cell division and differentiation areas of the C_4_ compared to C_3_ leaves, where vein formation takes place.

To test if developing *F. bidentis* leaves truly exhibit higher capacity of auxin homeostasis than the leaves of *F. robusta* we examined the resistance of both plant species towards externally applied synthetic auxins. Spraying the plants with NAA changed the overall leaf shape and significantly increased vascular density in *F. robusta* but did not affect leaves of *F. bidentis*. Application of 2,4-D on the other hand led to distorted leaf shape and significantly increased vascular density in both species in much lower concentrations (Figure 6). The higher sensitivity of both species toward 2,4-D compared to NAA is most likely due to the fact that NAA can be inactivated by GH3 proteins while 2,4-D, like other halogenated auxins, is not a substrate of these proteins (Staswick et al., 2005). This indicates that capacity for auxin homeostasis is indeed higher in the C_4_ species, while its obvious sensitivity towards 2,4-D confirms an involvement of GH3 proteins.

Veins form from procambial strands. These are induced by elevated auxin concentration due to polar auxin transport along strands of undifferentiated meristematic cells (Scarpella et al., 2004). According to the auxin canalization model (Sachs, 1969; Mitchison, 1981; Rolland-Lagan and Prusinkiewicz, 2005), such strands can form by local auxin maxima and induction of polar auxin transport due to PIN proteins. When more auxin is available in the cell division and differentiation areas of *F. bidentis*, more local auxin maxima could form and lead to the formation of procambial strands and, in the following, to the formation of more veins than in the C_3_ species *F. robusta*. This is in line with the observations of McKown and Dengler, (2009). They reported that the existence of an additional minor vein order (seven vein orders in *F. bidentis* compared to six vein orders in *F. robusta*) is the main reason for higher vein density in the C_4_ species. They observed an accelerated vein formation during C_4_ leaf development and concluded that this could be explained by either increased auxin production, modified leaf ground meristem cell competency to becoming procambium, or a combination of these developmental parameters (McKown and Dengler, 2009). According to our data, the C_4_ *Flaveria* can likely produce and process more auxin. Due to its enhanced auxin homeostasis capacity, it also can maintain higher auxin availability in the early cell differentiation area, where the higher order minor veins are induced (Figure 7). Hence, of the auxin x time = vein density equation, in *F. bidentis* the auxin term is modified, leading to higher vein density.

**Figure 7.**
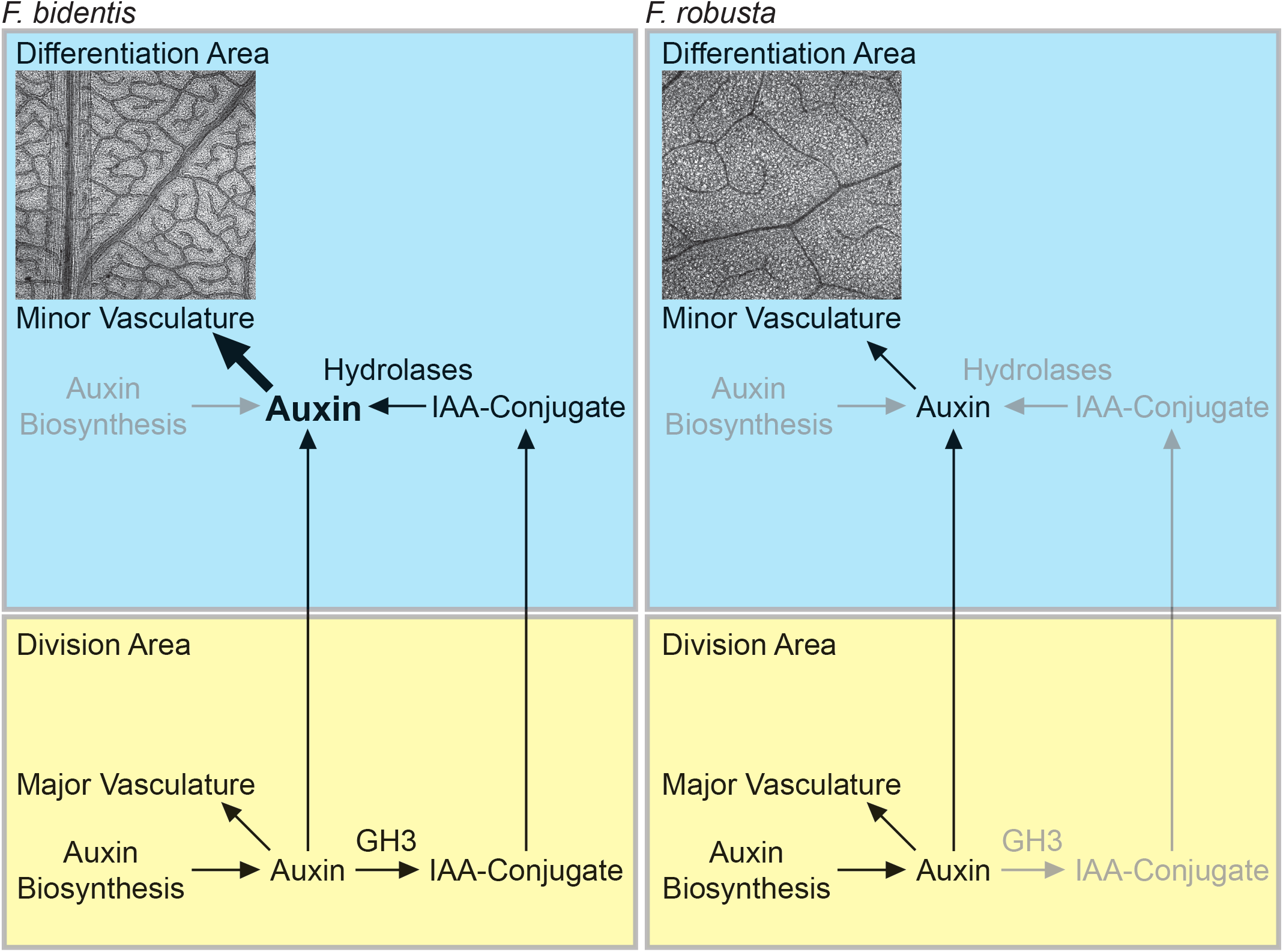
Differences in the induction of vein formation during leaf development in C_3_ and C_4_ *Flaveria* species.

### Polyphyletic evolution of C_4_ - comparison of *Flaveria* to other C_4_ origins

Given that C_4_ photosynthesis is a prime example of convergent evolution one might ask if the differences in leaf development we observed for C_3_ and C_4_ *Flaveria* species are specific for *Flaveria* only or can be generalized to other C_4_ species.

The interplay of the transcription factors SCARECROW (SCR) and SHORT-ROOT (SHR) is important for BS differentiation and to specify BS identity in C_3_ and C_4_ plants (Slewinski et al., 2012; Cui et al., 2014; Slewinski et al., 2014). Together with several IDD transcription factors these genes were found to be upregulated in dividing and differentiating tissues of the C_4_ leaves compared to the C_3_ leaves of the two *Flaveria* species (Supplemental Figure 10). Since these factors were also identified as candidate for BS differentiation in maize (Fouracre et al., 2014), one may assume that they represent common factors for the establishment of Kranz anatomy. They likely fulfill the same roles during C_4_ leaf development in multiple species and are found to be upregulated in developing C_4_ leaves because C_4_ leaves contain more BS cells compared to C_3_ leaves. The situation might be similar for factors involved in chloroplast development and maturation. We identified several factors like *GLK-2* or *CGA1* genes with different expression patterns in the developing leaves of the two *Flaveria* species. These are known to be essential for chloroplast development in C_4_ as well as in C_3_ plants (Rossini et al., 2001; Hall et al., 1998; Chiang et al., 2012; Wang et al., 2013a). When overexpressed in the BS cells of the C_3_ plant rice GLK genes led to C_4_-like differentiation of the rice BS (Wang et al., 2017b). In these cases, evolution of the C_4_ state converges on the same genes.

Regarding vein density, the model auxin x time = vein density provides at least two paths towards high vein density, either increasing the time for auxin exposure, or increasing the amount of available auxin. In *Flaveria* a combination of enhanced auxin synthesis and high auxin homeostasis capacity seem to accelerate vein formation by enhancing the induction of procambial strains in competent leaf areas. In contrast, for C_3_ and C_4_ species of Cleomaceae it was shown that high vein density is likely related to a delay in mesophyll cell differentiation during leaf development in the C_4_ species *Gynandropsis gynandra* (Külahoglu et al., 2014), probably in combination with elevated auxin synthesis (Huang et al., 2017). This leads to induction of more procambial strands and following more veins compared to the C_3_ species *Tarenaya hassleriana*, exhibiting a faster mesophyll differentiation (Külahoglu et al., 2014). Overall it appears that vascular density can be represented as a function of auxin concentration, auxin sensitivity and the duration of an inducible/competent state. In this framework, *Cleome* increases vascular density by prolonging the inducible state, while *Flaveria* keeps the length of the inducible state identical but increases auxin availability in tissues with the competence to differentiate into veins.

### Lessons for altering venation patterns prior to converting a C_3_ to a C_4_ species

There has been much interest in engineering C_3_ plants like rice to express C_4_ traits increasing their photosynthetic efficiency and productivity (Hibberd et al., 2008; Schuler et al., 2016). Leaf venation on its own is a trait predicted to influence photosynthetic capacity (Sack and Scoffoni, 2013). A critical step in engineering a C_4_ plant will be to increase leaf vein density. Obviously, parts of the equation auxin x time = vein density need to be altered to achieve altered leaf anatomy. Our analysis as well as earlier work (Külahoglu et al., 2014; Huang et al., 2017) demonstrates that different possibilities to alter auxin effects exist. It could be either a combination of enhanced auxin synthesis and prolongation of the time auxin is effective in vein initiation, as found in *Cleome*, or a combination of enhanced auxin synthesis and alterations in auxin homeostasis as we found for *Flaveria*. The most suitable way of alteration for engineering approaches has to be elucidated in future research.

## Materials and Methods

### Leaf material and RNA isolation

*Flaveria bidentis* and *Flaveria robusta* plants were grown from mid-October to mid-December in rooftop greenhouses at Düsseldorf university with at 16 h additional light per day, photon flux density (PFD) of ~300 μmol m^−2^ s^−1^ and at 24°C (during the day) and 21°C (at night). Leaf primordia along with shoot apical meristem (stage 0) and 9 leaves starting from first visible leaf pair to fully developed leaf pairs were harvested during noon and immediately frozen in liquid nitrogen. For each of the three biological replicates used in RNA-Seq analysis, leaf material was pooled from five plants and primordia (or shoot apices) were pooled from 25 plants. Approximate length of the different leaf stages was 0.3, 0.5, 1.5, 2, 3, 4, 6, 8 and 10 cm. Total RNA was isolated using the RNeasy Plant Mini Kit (QIAGEN) and DNase digestion was performed with RNase-Free DNase Set (QIAGEN). RNA Integrity Number (RIN), and quantity were determined with the 2100 Bioanalyzer (Agilent Technologies).

### Library preparation,sequencing and mapping

Libraries were prepared with the TruSeq RNA Library Prep Kit v2 (Illumina) using 0.5 μg of total RNA as a starting material. Quality and quantity of the libraries were determined with the 2100 Bioanalyzer and a Qubit instrument (Thermo Fisher Scientific), respectively, according to the instructions given by the manufacturer. Single end reads of 100 bp length were sequenced with the HiSeq2000 Illumina platform and samples were multiplexed with 6 libraries per lane. On an average ~ 33 million reads were obtained per sample. Reads were mapped to a minimal set of coding sequences of the *Arabidopsis thaliana* (http://www.Arabidopsis.org/), as described by Bräutigam et al., (2011). Read count was normalized to reads per kilobase of transcript, per million mapped reads (RPKM).

### RNAseq analysis and factorization

Unless stated otherwise all data analysis was performed with the R statistical package (version 3.3.3 “Another Canoe” www.r-project.org). The transcriptome was filtered by expression and all genes with an expression above ten RPKM in at least one sample across both species were considered for further analysis. Transcriptional investment was calculated based on reduced Mapman categories (Brilhaus et al., 2016; Külahoglu et al., 2014). PCA analysis was performed on Z-scored expression values.

We factorized our data using altered variants of the Lee and Suad, the Brunet, the Kullback-Leibler and the non-smooth NMF algorithms provided within the NMF package for R (Gaujoux and Seoighe, 2010). The used factorizations were performed by iterative matrix multiplications based on euclidean distance in the Lee an Seung algorithm and Kullback Leibler divergence in the Brunet and KL algorithms. Non smooth NMF (nsNMF) additionally imposes sparseness on the resulting factorization by incorporating a smoothing matrix into every iteration (Pascual-Montano et al., 2006). Different sparsities were tested by using theta values of 0.3, 0.5, 0.7 and 0.9 with 0.3 creating the densest and 0.9 creating the sparsest result. In general, NMF is an ill composed task with several possible solutions to a factorization, represented by local minima in the distance-function. Therefore, factorization was performed several times with different starting points. To this end we utilized three different seeding methods, random seeding, independent component analysis based seeding (ica) and Nonnegative Double Singular Value Decomposition (nndsvd) seeding (Boutsidis and Gallopoulos, 2008). Due to its computational costs we performed the initial factorizations determining the rank of the NMF and comparing different methods and initializations fifty times for two to ten areas (Supplementary Figure 2). To improve calculation speed, factorization was performed on the transpose of the Gene by Sample matrix. The coefficient matrix was restricted to represent tissue structure by dividing each value in a column with its sum during every iteration. The final factorizations were performed 500 times using the Kullback-Leibler algorithm for five areas, when the primordia were included and for four areas for the dataset without primordia.

We clustered the factorized data using the K-means algorithm (Hartigan and Wong, 1979) after Z-scoring by genes across both species. A suitable number of clusters was determined using the elbow method and clustering was performed 2000 times. GO enrichment was performed using the TopGo package in R (Alexa and Rahnenführer, 2018). GO annotations were taken from TAIR10 and the enrichment was detected by Fisher’s Exact Test. All generated p-values were corrected for multiple testing using Benjamini-Hochberg correction (Benjamini and Yosef, 1995). The comparative PCA (Figure 3B) was performed on the data provided in the supplementary dataset of Kümpers et al., (2017). Genes were filtered for AGIs present in both studies and by gene normalization was performed separately for the Kümpers dataset, the initial dataset and the factorized dataset.

### Araldite embedding

For Araldite embedding we cut 1 x 2 mm sized segments from the second quarter of the leaves. The samples were transferred into 4% paraformaldehyde solution until the majority of the segments dropped to the bottom of the tube or for at least 24h. The material was transferred into 0.1% v/v glutaraldehyde in 1x PBS, incubated for 20 min at room temperature and vacuum infiltrated three times for five minutes each. Subsequently an ethanol dilution series from 30% to 90% ethanol, with an increment of 10% for each step was applied. Each step was carried out for 20 minutes and the final dehydration steps were carried out in 96% ethanol (two times), 100% ethanol (two times) and 100% acetone (two times) for one hour. To remove residual water the 100% ethanol and acetone solutions were dried using zeolite capsules. Finally, the samples were transferred into open vessels and overlaid with araldite. To ensure acetone evaporation the samples were incubated for 4 hours at room temperature under a fume hood. The samples were transferred into fresh araldite in a silicon mold and incubated overnight. The final polymerization was performed for 24-48h at 65°C. Samples were cut with a microtome, stained in bromophenol blue, and observed at 20-fold magnification.

### Leaf clearing and determination of vascular structure

Leaf clearing was performed as described by Hasegawa et al., (2016). Leaves of a suitable size were cleared in ethanol acetic acid (3:1 v/v) until no green color was visible and transferred to 97% 2,2’-thiodiethanol solution containing 0.0025% (w/v) propyl gallate in PBS for 20 minutes. All pictures were taken at four times magnification with an estimated area overlap of 30 to 50% and subsequently stitched together in Adobe Photoshop (Version 2014.0.0), using the build in merge function. The vascular density estimation was performed in 200 μm steps for leaves up to 4 cm length and in 500 μm steps for larger leaves.

### Auxin treatment

The effect of auxin on leaf structure and vascular density of *F. robusta* and *F. bidentis* was tested by spraying with NAA and 2,4-D. *Flaveria* seeds were surface sterilized, germinated for two weeks on half strength MS medium with 0.8% (w/v) agar in a 16 h light/ 8 h dark cycle (22/18°C) and transferred to soil. One week after the transfer, plants were sprayed for three weeks on a daily basis with either 1-Naphthaleneacetic acid (NAA, Sigma Aldrich) or 2,4- Dichlorophenoxyacetic acid (2,4-D, PESTANAL Sigma Aldrich). Both phytohormones were dissolved in DMSO and 0.1% v/v DMSO was used as negative control.

### Accession numbers

The read data have been submitted to the National Center for Biotechnology Information Short Read Archive under accession numbers: XXXX (*F. bidentis*) and YYYY (*F. robusta*)

## Acknowledgements

This work was supported by the Deutsche Forschungsgemeinschaft through the Excellence Cluster EXC 1028 (From Complex Traits towards Synthetic Modules) and the International Graduate Training Program iGRAD-Plant (IRTG 1525). We thank the “Genomics and Transcriptomics laboratory” of the “Biologisch-Medizinischen Forschungszentrum” (BMFZ) at the Heinrich-Heine-University of Düsseldorf (Germany) for technical support and conducting the Illumina sequencing.

## Authors contributions

Conception and design: KB, TW, AB, APMW, PW, UG; Acquisition of data: KB, TW; Analysis and interpretation of data KB, TW, AB, UG; Drafting or revising the article: KB, TW, AB, APMW, PW, UG;

## Supplemental Material

**Supplemental Table 1** Results of the Illumina sequencing runs and read mapping.

**Supplemental Table 2** CSV table providing quantitative information for all reads mapped onto the reference transcriptome from *Arabidopsis thaliana*

**Supplemental Table 3** CSV table providing quantitative information for all reads mapped onto the reference transcriptome from *Arabidopsis thaliana* targeting genes expressed with at least ten RPKM used for further analysis, read counts normalized to RPKM, modeled RPKM data, average regression coefficients and cluster affiliations

**Supplemental Table 4** CSV table providing the results of GO enrichment analysis of transformed and clustered expression data including the leaf primordia data

**Supplemental Table 5** CSV table providing the results of GO enrichment analysis of transformed and clustered expression data without the leaf primordia data

**Supplemental Figure 1** Cross sections of the developmental progression in *F. bidentis* and *F. robusta*

Cross sections were taken in the third quarter of the leaf for every step in the developmental gradient. The numbers represent the gradient stage with the corresponding leaf lengths of 0.5 cm, 1.5 cm, 2.0 cm, 3.0 cm, 4.0 cm, 6.0 cm, 8.0 cm, and 10.0 cm. The *F. bidentis* section is depicted at the top and *F. robusta* at the bottom of each pair. Scale bar – 10 μm.

**Supplemental Figure 2** Determination of the parameters for nonnegative matrix factorization The quality of NMF factorizations for two to ten factors was determined to test different NMF algorithms (A) and different seeding methods (B). The suitable number of areas was determined via the residual sum of squares (C) and the average regression coefficient (D).

**Supplemental Figure 3** Factorization of the dataset with primordia

The dataset containing the primordia samples was factorized into five areas. The relative proportions of an area contributing to the sample set for both species is presented and the samples sorted according to their corresponding leaf length with an assigned length of 0 cm for primordia.

**Supplemental Figure 4** K-means clusters based on factorized gene expression for the dataset including primordia

The dataset containing the primordia samples was clustered into seven clusters based on the factorized expression profiles within the five areas. GO terms overrepresented in both species in clusters of similar progression.

**Supplemental Figure 5** PCA of the leaf developmental gradient

PCA of the developmental gradient for all leaf stages sequenced in this study (circles) as well as the *Flaveria* slice transcriptomes from Kümpers *et al*. 2017 (diamonds). The modelled areas are depicted in green (*F. bidentis*) and black (*F. robusta*) and represented by squares.

**Supplemental Figure 6** K-means clusters based on factorized gene expression for the dataset without primordia

The dataset without the primordia samples was factorized into four areas and k-means clustered based on the intensity of the area specific expressions into seven clusters.

**Supplemental Figure 7** Heatmap of genes related to the C_4_ cycle

Gene expression is presented normalized separately for the modelled and the sampled data to the respective maximum in both species.

**Supplemental Figure 8** Heatmap of genes related to Cyclic electron transport

Gene expression is presented normalized separately for the modelled and the sampled data to the respective maximum in both species.

**Supplemental Figure 9** Heatmap of genes related to Photorespiration

Gene expression is presented normalized separately for the modelled and the sampled data to the respective maximum in both species.

**Supplemental Figure 10** Heatmap of genes related to Bundle sheath differentiation

Gene expression is presented normalized separately for the modelled and the sampled data to the respective maximum in both species.

**Supplemental Figure 11** Heatmap of genes related to Auxin signalling

Gene expression is presented normalized separately for the modelled and the sampled data to the respective maximum in both species.

**Supplemental Figure 12** Heatmap of Auxin induced genes

Gene expression is presented normalized separately for the modelled and the sampled data to the respective maximum in both species.

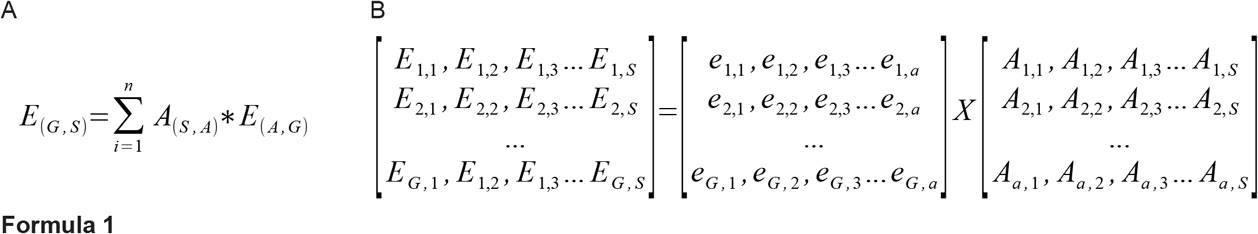

